# NKX2.1 is critical for melanocortin neuron identity, hypothalamic *Pomc* expression and body weight

**DOI:** 10.1101/460501

**Authors:** Daniela P. Orquera, M. Belén Tavella, Flavio S. J. de Souza, Sofía Nasif, Malcolm J. Low, Marcelo Rubinstein

**Author notes:** These authors contributed equally (Daniela P. Orquera, M. Belén Tavella). Correspondence to: INGEBI-CONICET, Vuelta de Obligado 2490, 1428-Buenos Aires, ARGENTINA.

## Abstract

Food intake is tightly regulated by a group of neurons present in the mediobasal hypothalamus which activate satiety by releasing *Pomc*-encoded melanocortins. Although the relevance of hypothalamic POMC neurons in the regulation of energy balance and body weight is well appreciated, little is known about the transcription factors that establish their cellular fate, terminal differentiation and phenotypic maintenance. Here, we report that the transcription factor Nkx2.1 activates hypothalamic *Pomc* expression from early development to adulthood by binding to conserved canonical NKX motifs present in the neuronal *Pomc* enhancers nPE1 and nPE2. Transgenic and mutant mouse studies showed that the NKX motifs present in nPE1 and nPE2 are essential for their transcriptional enhancer activity. Early inactivation of *Nkx2.1* in the ventral hypothalamus prevented the onset of *Pomc* expression and selective *Nkx2.1* ablation from POMC neurons impaired *Pomc* expression and increased body weight and adiposity. These results demonstrate that NKX2.1 is critical in the early establishment of arcuate melanocortin neurons and the regulation of *Pomc* expression and body weight in adulthood.

---

The arcuate nucleus of the hypothalamus is a brain hub that integrates nutritional and hormonal information to promote food intake or satiety (1). A group of arcuate neurons expresses proopiomelanocortin (*Pomc*), a gene that encodes the anorexigenic neuropeptides α-, β- and γ-melanocyte-stimulating hormone (MSH), collectively known as central melanocortins (1). POMC neurons promote satiety upon sensing variations in glucose (2), leptin (3) and insulin (4) levels as well as in local temperature (5). In addition, POMC neurons receive multiple synaptic connections from a variety of neurons that, together, orchestrate food intake (6-10). The physiological importance of central melanocortins is evident in hypothalamic *Pomc*-deficient mice, which are hyperphagic and display early onset extreme obesity (11,12). In addition, humans (13) and mice (14) carrying null allele mutations in *POMC* or in the melanocortin receptor 4 (15,16) are also hyperphagic and severely obese. Although POMC neurons play a fundamental role in the regulation of food intake and body weight, little is known about the genetic programs that establish their identity and phenotypic maintenance.

Hypothalamic *Pomc* expression is controlled by two upstream distal enhancers, nPE1 and nPE2, which are highly conserved in mammals (17). Although nPE1 and nPE2 have unrelated evolutionary origins, both enhancers drive overlapping spatiotemporal activities to the entire population of hypothalamic POMC neurons (18,19). Targeted mutagenesis of nPE1 and/or nPE2 revealed their partially redundant enhancer function and cooperativity to maintain *Pomc* levels above a critical functional threshold (20). Interestingly, only the concurrent removal of both enhancers reduced *Pomc* expression to very low levels, leading to hyperphagia and early-onset obesity (20). Because nPE1 and nPE2 act as transcriptional *Pomc* enhancers in the same population of hypothalamic neurons, it is conceivable that they share DNA elements that recruit similar transcription factors (TF). In fact, we detected a 21-bp sequence embedded in a highly conserved region of nPE1 and a similar sequence in nPE2 (20) which contains ATTA motifs typically recognized by homeodomain TFs (20). Recently, we found that these motifs are recognized by Islet 1 (ISL1), a LIM-homeodomain TF that coexpresses with *Pomc* in the developing hypothalamus and postnatal life (21). Moreover, early inactivation of *Isl1* prevents the onset of hypothalamic *Pomc* expression, and ablation of *Isl1* from POMC neurons impairs *Pomc* expression in adult mice and leads to hyperphagia and obesity (21).

Although ISL1 is necessary for *Pomc* expression, it is clearly not sufficient. *Isl1* is expressed not only in a much broader hypothalamic territory than *Pomc*, but also in other neuronal types where *Pomc* is never found (22-24). Thus, other TFs are likely to participate in the arcuate-specific expression of *Pomc*. Here, we combined molecular, cellular, genetic and functional studies we show to demonstrate that the homeodomain TF NKX2.1 plays a crucial role in arcuate *Pomc* expression from early development to adulthood by interacting with motifs present in nPE1 and nPE2. Therefore, NKX2.1 defines the identity of hypothalamic melanocortin neurons and is necessary to maintain sufficient levels of hypothalamic *Pomc* mRNA in adult mice to assure normal body weight regulation.

## Results

### NKX2.1 is a candidate TF for the regulation of hypothalamic *Pomc* expression

Within the neuronal *Pomc* enhancer nPE1 we detected a canonical binding motif for a TF of the NKX subfamily which is highly conserved in all mammalian orders (Fig. 1, left and Fig. S1) while two conserved canonical NKX binding sites are present in nPE2 (Fig. 1, right and Fig. S2). Among all NKX family members, NKX2.1 emerged as the most likely candidate to regulate hypothalamic *Pomc* expression because it is present in the early developing ventral hypothalamus portion that gives rise to the arcuate nucleus (25–28), and continues to be expressed in this nucleus during postnatal life (29). A necessary condition to support NKX2.1 as a transactivator of hypothalamic *Pomc* is that both genes coexpress within the same neurons. We followed the developmental pattern of *Nkx2.1* expression in the ventromedial hypothalamus in sagittal sections of *Pomc*-EGFP mice, a validated transgenic line that coexpresses EGFP and *Pomc* along the entire spatiotemporal domain and that is useful to maximize the identification of POMC neurons (Fig. S3) (3, 21). At the onset of *Pomc* expression in the mouse hypothalamus (E10.5) (30), all incipient *Pomc*-EGFP neurons also express *Nkx2.1* (Fig. 2 a,b,c). The expression territory of *Nkx2.1* is much broader than that of *Pomc* extending beyond the limits of the future arcuate nucleus. Expression of *Nkx2.1* in all POMC neurons is also evident at E12.5, when neurogenesis of POMC neurons is highest (Fig. 2 d,e,f). In the adult arcuate nucleus, most *Pomc*-EGFP positive-neurons are immunopositive for NKX2.1 (Fig. 2 g,h,i) but, as in earlier stages, the total number of NKX2.1 immunopositive neurons is greater than that of POMC neurons. Together, these results show that POMC neurons express *Nxk2.1* in the ventromedial hypothalamus during development and adulthood.

**Figure 1.**
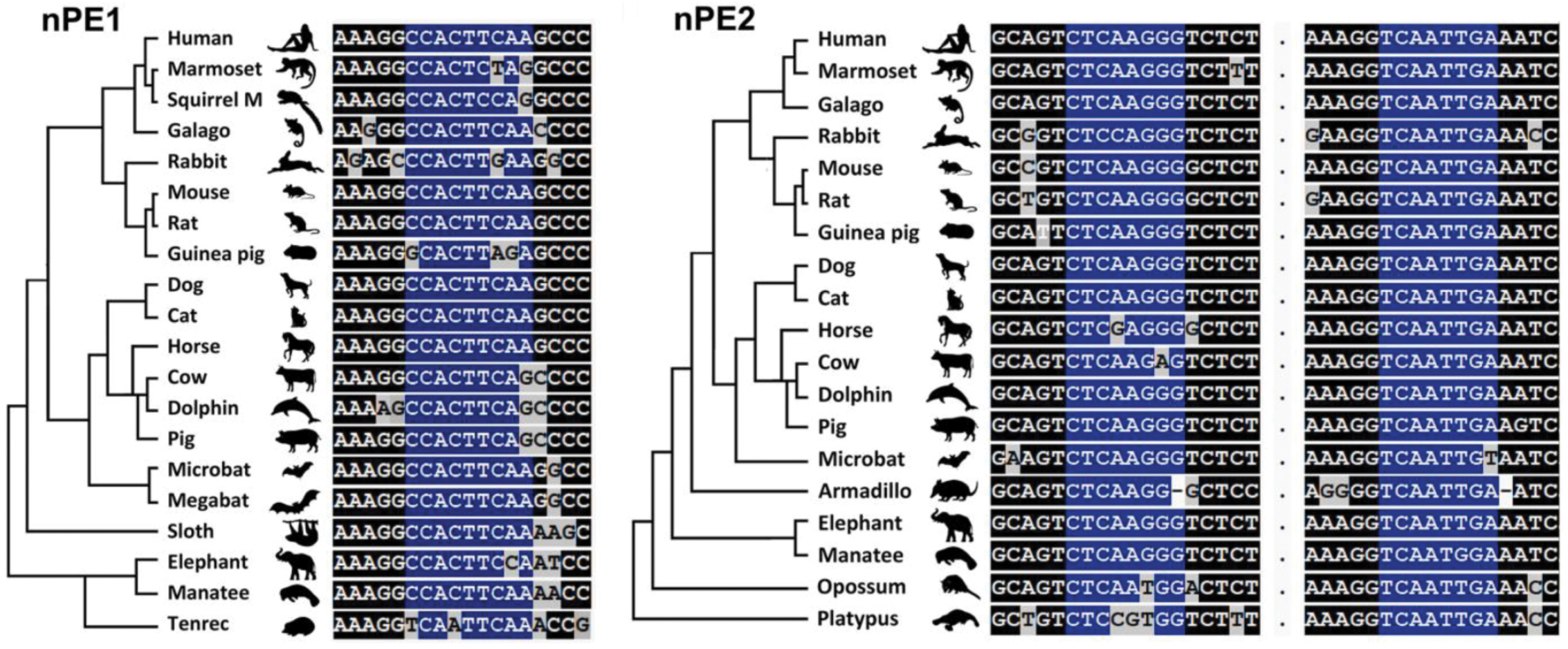
The *Pomc* neuronal enhancers nPE1 and nPE2 contain canonical binding motifs for a transcription factor of the NKX family which are highly conserved in all mammalian orders. (Left) Evolutionary tree of placental mammals followed by a ClustalW alignment showing the unique NKX binding site in nPE1. (Right) Evolutionary tree of mammals followed by a ClustalW alignment of the two NKX binding sites present in nPE2. Conserved nucleotides are indicated in a dark background and those conserved nucleotides corresponding to canonical NKX binding sites in blue.

**Figure 2.**
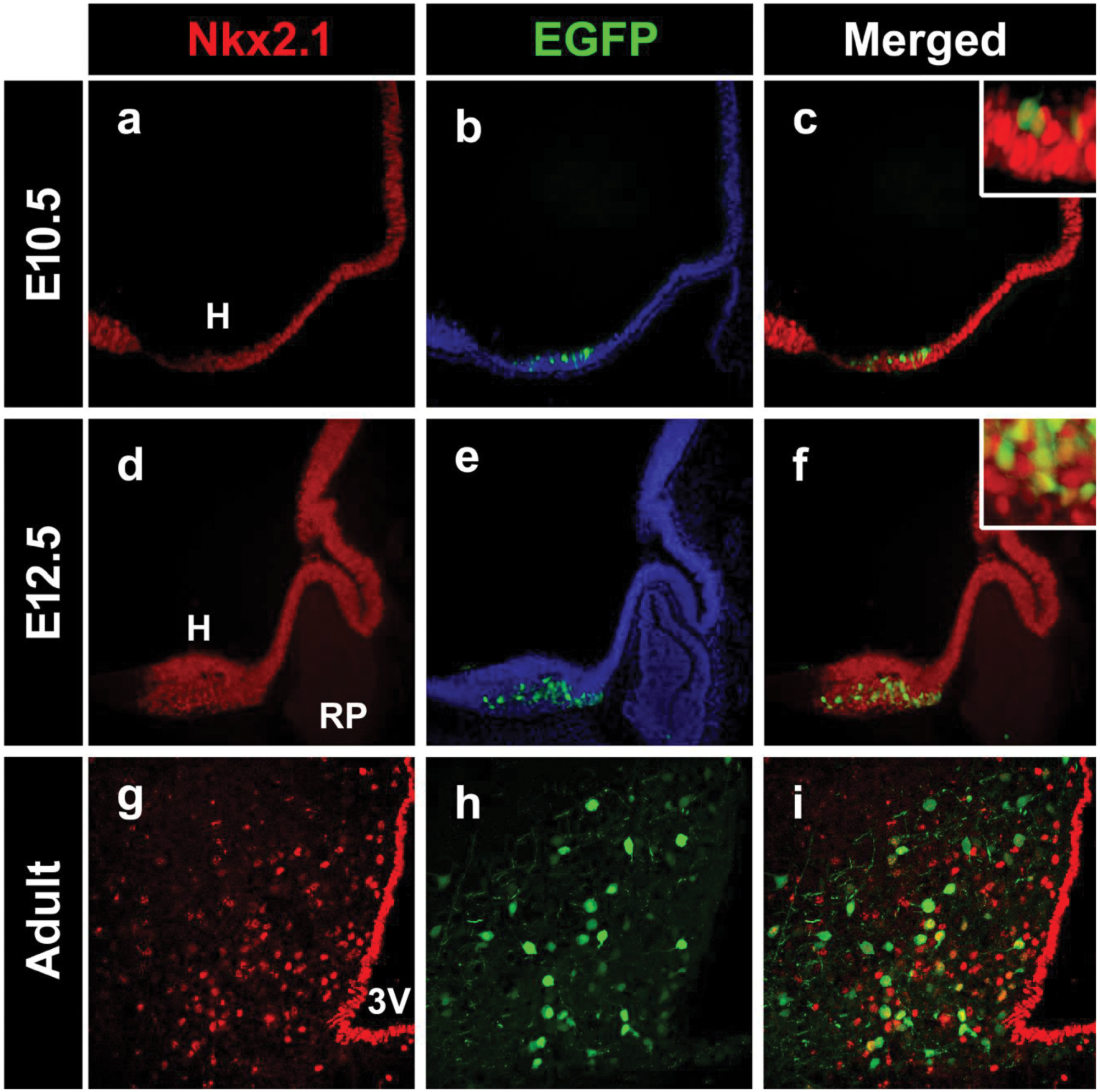
*Pomc* coexpresses with *Nkx2.1* in the developing and adult mouse hypothalamus. Expression analysis of *Nkx2.1* in sagittal cryosections of *Pomc*-EGFP mouse E10.5 (a-c) and E12.5 (d-f) embryos and coronal sections of adult *Pomc*-EGFP mice at the level of the arcuate nucleus (g-i). Insets are magnified views showing that all EGFP+ cells express *Nkx2.1*. H, future hypothalamus; RP, future Rathke’s pouch; 3V, third ventricle.

### NKX2.1 specifically binds *in vitro* and *in vivo* to DNA elements present in nPE1 and nPE2

We then examined the *in vitro* binding properties of NKX2.1 in electrophoretic mobility shift assays (EMSA) using a bacterially expressed mouse *Nkx2.1* clone. We found that 30-bp DNA probes encompassing canonical NKX motifs present in nPE1 or nPE2 were shifted when incubated with NKX2.1 bacterial extracts (Fig. 3a). To test whether NKX2.1 interacts with nPE sequences *in vivo* we performed a chromatin immunoprecipitation (ChIP) assay using chromatin harvested from adult mouse hypothalami. An anti-NKX2.1 antibody pulled down nPE1 and nPE2 sequences that were amplified by PCR whereas a control IgG antibody failed to immunoprecipitate either nPE sequence (Fig. 3b,c).

**Figure 3.**
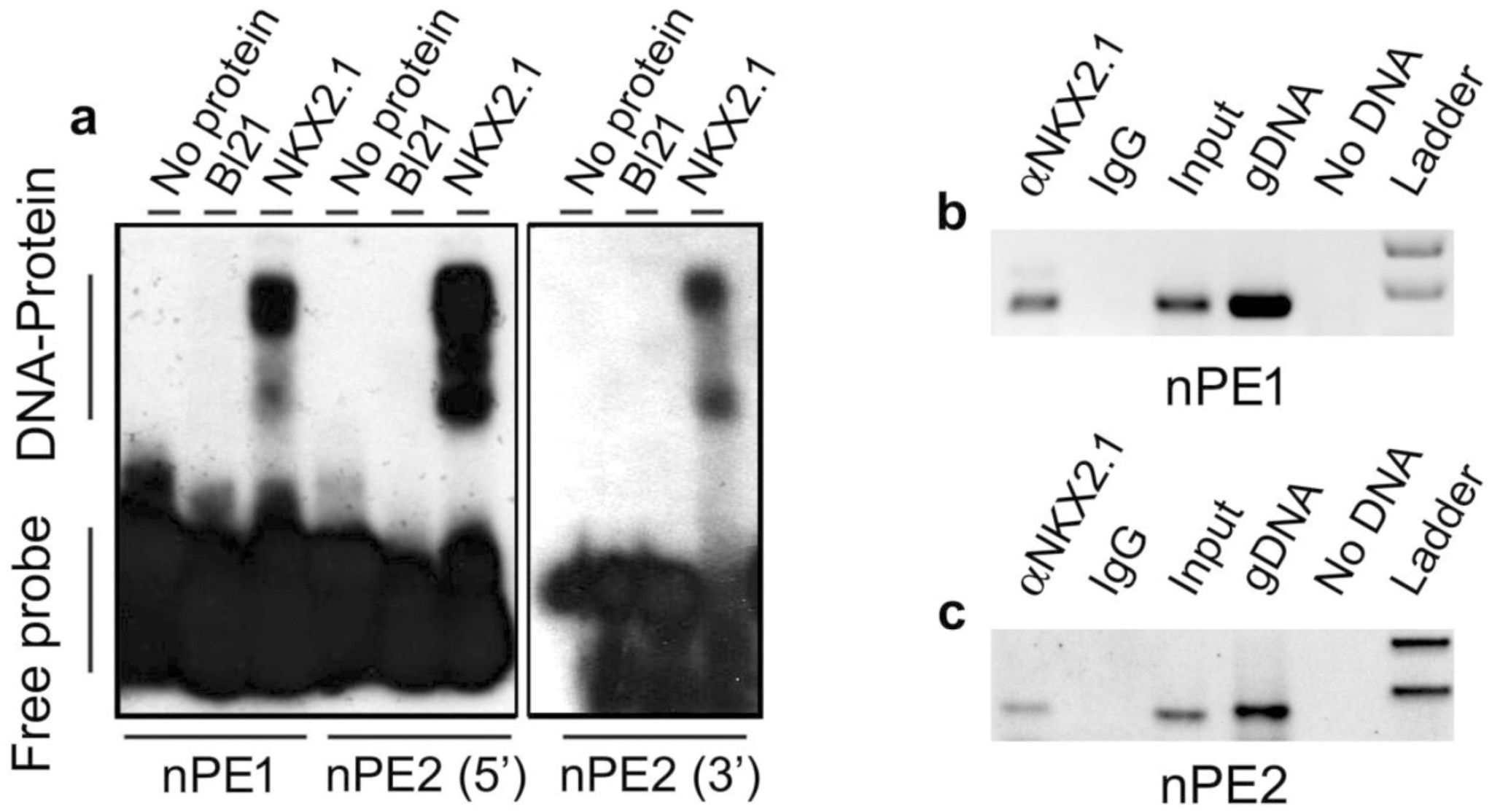
NKX2.1 binds to nPE1 and nPE2 sequences in vitro and in vivo. (a) EMSA shows that NK2.1 binds to the NKX canonical sequence present in nPE1 (left) and to two NKX motifs present in nPE2 (5’site middle, 3’ site right). (b, c) ChIP assay with adult mouse hypothalamus chromatin and an anti-NKX2.1 antibody that immunoprecipitates nPE1 (b) and nPE2 (c) sequences.

### Functional analysis of NKX binding sites present in the neuronal *Pomc* enhancers nPE1 and nPE2

In previous studies we demonstrated that the sole presence of nPE1 or nPE2 in transgenic constructs is sufficient to drive reporter gene expression to arcuate POMC neurons (17,19). To determine the functional relevance of the two NKX binding motifs present in nPE2 (Fig. 1b), we tested whether a mutant version carrying transition mutations in these two binding sites is able to drive EGFP expression to the arcuate nucleus of transgenic mice (Fig. 4a). Both transgenes carried the mouse *Pomc* proximal promoter known to drive reporter gene expression exclusively to POMC pituitary cells which allows us to discard transgenic mice carrying silent integrations (18, 31). As expected from previous studies (16), we found that transgene nPE2*Pomc*-EGFP drove expression to arcuate hypothalamic neurons in three different pedigrees (Fig. 4b). In contrast, four independent transgenic lines carrying the mutant version nPE2(NKX*)*Pomc*-EGFP displayed either very few (two lines, Fig. 4c) or complete absence of EGFP positive-arcuate neurons (two lines, Fig. 4d). All seven transgenic lines analyzed displayed consistent EGFP expression in the pituitary gland (Fig. 4e-g). These results indicate that the NKX binding motifs in nPE2 are critical for proper enhancer activity in arcuate POMC neurons.

**Figure 4.**
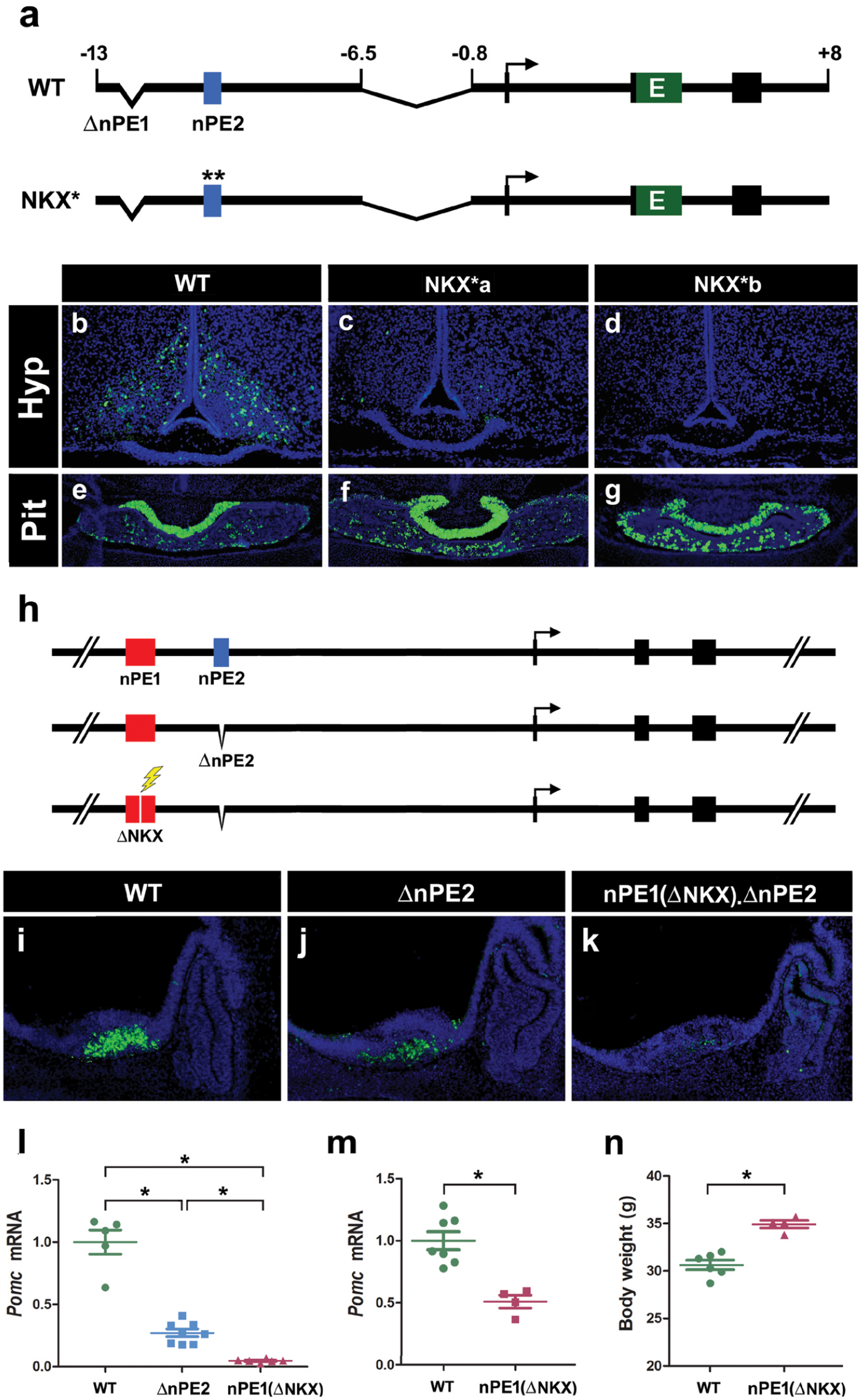
NKX2.1 binding sites in nPE1 and nPE2 are important for enhancer function. (a) Schematic of two nearly identical transgenes constructed to study the importance of the NKX binding motifs present in nPE2. Asterisks indicate the transversion mutations (AA→CC) introduced in the two NKX binding sites present in nPE2 (NKX*). (b-g) Expression analysis of transgenic mice at P0 carrying wild-type nPE2 (WT) or the mutant nPE2 version driving EGFP on coronal sections at the level of the arcuate nucleus (b-d) and the pituitary (e-g). Two independent transgenic lines carrying nPE2(NKX*) are shown (c,d,f,g). (h) Schematic of the three *Pomc* alleles analyzed at E12.5. WT mice carry intact nPE1 and nPE2, ΔnPE2 mice lack nPE2, and nPE1(ΔNKX). ΔnPE2 mice also carry a deletion of the NKX binding motif in nPE1 introduced by CRISPR/Cas9, indicated with a lightning symbol. (i-k) Immunofluorescence analysis of *Pomc* expression using an anti-ACTH antibody on sagittal sections in E12.5 embryos. (l) RT-qPCR analysis of *Pomc* mRNA levels collected from *Pomc^+/+^*, *Pomc^.ΔnPE2/ΔnPE2^* and *Pomc^nPE1(ΔNKX).ΔnPE2/ nPE1(ΔNKX).ΔnPE2^* E11.5 embryo heads. Bars represent the mean +/− SEM of 5-8 samples. One-way ANOVA followed by Tukey posthoc test,*p<0.001. (m) RT-qPCR analysis of hypothalamic *Pomc* mRNA levels and (n) Body weight from *Pomc+/+* and *Pomc^nPE1(ΔNKX).ΔnPE2/ nPE1(ΔNKX).ΔnPE2^* littermates at 12 week-old. Bars represent the mean +/− SEM of 4-6 samples. Student’s t test, *p<0.001.

To test the importance of the unique NKX binding site present in nPE1, we took a different genetic approach using CRISPR/Cas9 technology to selectively delete the motif from nPE1 by targeted mutagenesis of *Pomc* alleles already lacking nPE2 (20). We microinjected *Pomc^+/ΔnPE2^* zygotes with a sgRNA directed to the unique NKX site present in nPE1 and obtained nPE1(ΔNKX) mutant alleles that, in addition, lack nPE2 (*Pomc^nPE1(ΔNKX).ΔnPE2^*; Fig. 4h). *Pomc^ΔnPE2/ΔnPE2^* E12.5 embryos showed reduced *Pomc* expression levels compared to *Pomc^+/+^* controls, in agreement with our previous findings (Fig. 4i,j; 20). Interestingly, homozygous carriers of *Pomc^nPE1(ΔNKX).ΔnPE2^* alleles showed even greater reductions in *Pomc* expression levels in the developing arcuate nucleus compared to *Pomc ^ΔnPE2/ΔnPE2^* mutants, with only occasional ACTH+ neurons observed (Fig. 4i-k). A quantitative analysis performed by RT-qPCR revealed that homozygous *Pomc^ΔnPE2/ΔnPE2^* E11.5 embryos express 27.1 % (p<0.001) of wild-type *Pomc* mRNA levels (Fig. 4l), similar to what has been previously reported (20). Interestingly, ablation of the NKX binding motif present in nPE1 (*Pomc^nPE1(ΔNKX).ΔnPE2^*) further reduced *Pomc* expression to only 4.6 % (p<0.001) of normal levels (Fig. 4l) present in *wild-type* E11.5 siblings. Different to *Pomc^ΔnPE2/ΔnPE2^* adult mice which express 80% of hypothalamic *Pomc* mRNA levels and display normal body weight (20), homozygous carriers of *Pomc^nPE1(ΔNKX).ΔnPE2^* express only 51% of *Pomc* mRNA levels (Fig. 4m) and are significantly heavier (34.9 +/− 0.4 g) than their wild type littermates (30.6 +/− 0.5 g; Fig. 4n). These results reveal that the NKX motif present in nPE1 is a functionally critical component of this enhancer. Altogether, our transgenic and mutant studies demonstrate that the neuronal *Pomc* enhancers nPE1 and nPE2 depend on their NKX binding sites to fully drive endogenous or reporter gene expression in hypothalamic neurons and suggest that NKX2.1 is critically involved in neuronal *Pomc* expression.

### NKX2.1 is critical for the early establishment of hypothalamic melanocortin neuron identity

To investigate whether NKX2.1 participates in the establishment of the hypothalamic POMC lineage and/or in the developmental expression of *Pomc* we used a conditional mutant mouse strain that allowed *Nkx2.1* ablation at different developmental time points. In *Nkx2.1^loxP/loxP^* mice, the homeodomain-encoding exon 2 is flanked by two loxP sites so that null alleles are generated upon Cre recombinase activation (32). By crossing *Nkx2.1^loxP/loxP^* conditional mutants with mice harbouring a transgene that ubiquitously expresses a tamoxifen-inducible Cre recombinase (CAAG-CreERT) (33) we obtained inducible *Nkx2.1^loxP/loxP^*.CAAG-CreER mice, that for simplicity we named Ind*Nkx2.1*KO. Pregnant *Nkx2.1^loxP/loxP^* dams mated with Ind*Nkx2.1*KO males received a single tamoxifen (TAM) injection at different developmental time points (E8.5, E9.5 and E10.5) and Ind*Nkx2.1*KO and *Nkx2.1^loxP/loxP^* littermates were collected at E12.5, with the latter being used as controls. Ind*Nkx2.1*KO embryos receiving TAM at E8.5 (*Nkx2.1*KO@E8.5) showed complete absence of NKX2.1 when evaluated in E12.5 sagittal sections (Fig. 5 a,b), demonstrating that this dose of TAM induced efficient recombination of the conditional *Nkx2.1^loxP^* alleles. *Nkx2.1*KO@E8.5 embryos showed a thinning of the ventral neuroepithelium at the level of the future hypothalamus and lack of infundibulum, as found in *Nkx2.1-/-* mice (40). Immunofluorescence performed in *Nkx2.1*KO@E8.5 embryos showed only a few POMC immunoreactive cells, in contrast to what was observed in control embryos lacking CreER (Fig. 5 c,d). TAM injected at E9.5 also induced inactivation of the *Nkx2.1* conditional alleles as evidenced by the complete lack of NKX2.1 signal in *Nkx2.1*KO@E9.5 embryos that showed an overall normal morphology (Fig. 5 e,f). In the absence of Nkx2.1, the number of POMC positive-cells was greatly diminished in *Nkx2.1*KO@E9.5 embryos (Fig. 5 g,h). These results indicate that *Nkx2.1* ablation from early stages of embryonic development impairs the onset of *Pomc* expression. Thus, Nkx2.1 is necessary to fully activate hypothalamic *Pomc* expression and, consequently, to establish the entire set of melanocortin neurons.

**Figure 5.**
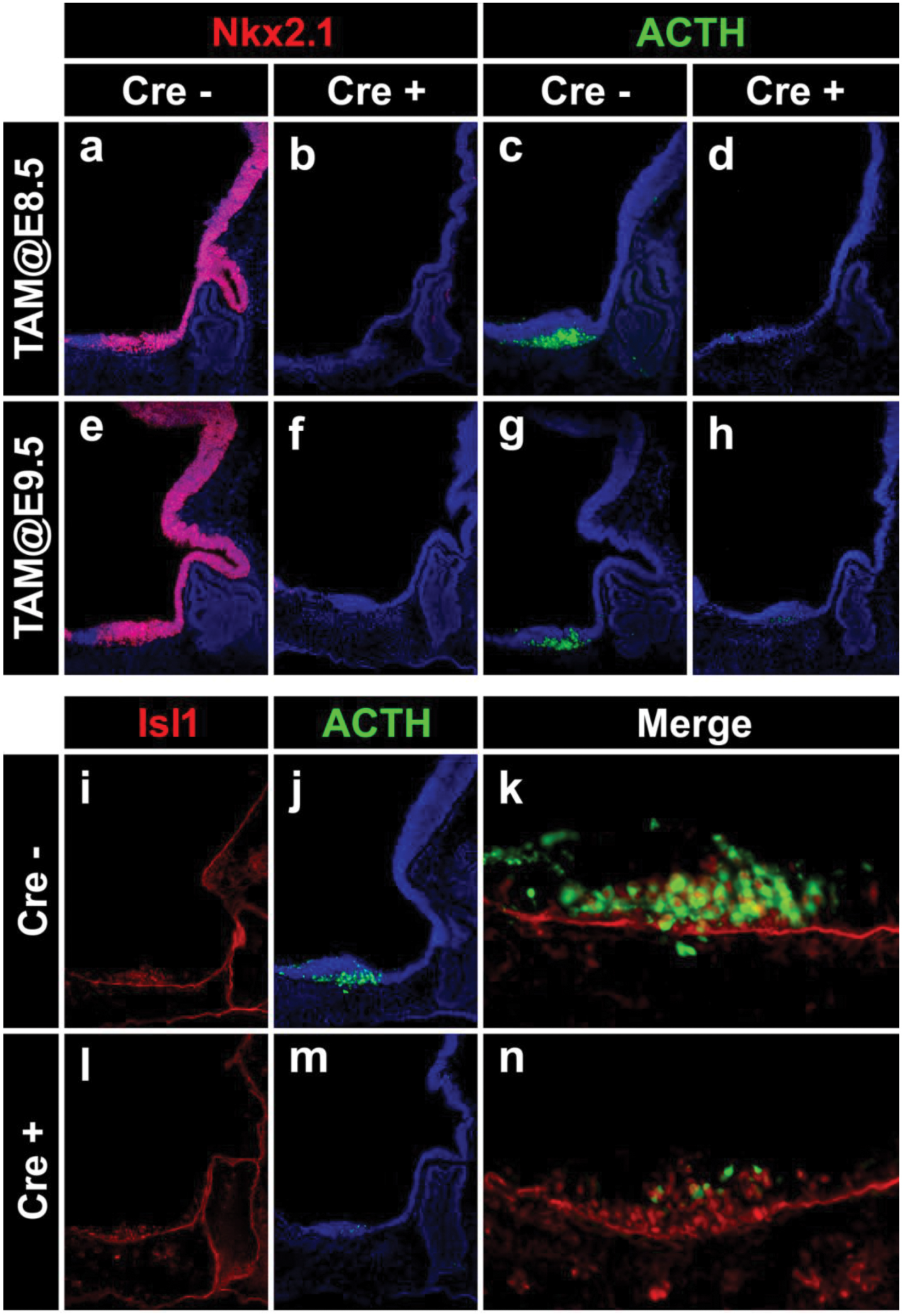
Hypothalamic *Pomc* expression is impaired in *Nkx2.1* conditional knockout mice. (a-h) Immunofluorescence analysis using anti-NKX2.1 (red) and anti-ACTH (green) antibodies in sagittal cryosections of E12.5 mouse embryos injected with tamoxifen at E8.5 (a-d) or E9.5 (e-h). Cre+ and Cre-embryos are littermates. (i-n) Immunofluorescence analysis of E12.5 mouse embryos injected with tamoxifen at E9.5 using antibodies against Isl1 (i,l) or ACTH (j,m). Close-up merged pictures are also shown (k,n).

In a previous study we showed that the onset of hypothalamic *Pomc* expression at E10.5 depends on the earlier expression of the TF ISL1, that starts at E10.0 in maturing postmitotic neurons of the future arcuate nucleus (21). Given that the onset of *Nkx2.1* expression in the developing hypothalamus precedes that of Isl1, we tested the hypothesis that NKX2.1 could also activate *Pomc* expression via ISL1 alternatively to its direct effect acting on the NKX binding sites present in nPE1 and nPE2. We found, however, normal ISL1 immunoreactivity in ventromedial hypothalamic neurons of *Nkx2.1*KO@E9.5 embryos (Fig. 5i, l), indicating that *Isl1* does not seem to be downstream of *Nkx2.1*. Furthermore, in the absence of Nkx2.1, Isl1 was unable to promote normal *Pomc* expression as evidenced by the large number of Isl1 immunoreactive neurons that failed to express *Pomc* in the hypothalamus of *Nkx2.1*KO@E9.5 embryos (Fig. 5l-n), in comparison to what was found in their *Nkx2.1^loxP/loxP^* siblings (Fig. 5i-k). These results, together with those shown in the sections above, support the idea that NKX2.1 activates *Pomc* expression directly.

### *Nkx2.1* expression in ISL1 neurons is critical for hypothalamic *Pomc* expression

To limit *Nkx2.1* ablation to the lineage leading to arcuate POMC neurons we decided to inactivate *Nkx2.1* alleles specifically in neurons expressing *Isl1*. We found that the pattern of *Isl1* expression within the presumptive arcuate nucleus at E10.5 overlaps with that of *Nkx2.1* (Fig. 6 a,b,d), and that POMC positive-cells coexpress *Isl1* and *Nkx2.1* (Fig. 6 c,e). A similar pattern of triple coexpression was observed in E12.5 embryos, in which more than 80% of POMC neurons coexpressed both TFs (Fig. 6 f-j). By mating *Nkx2.1^loxP/loxP^* mice with a knockin strain that expresses Cre under the transcriptional control of *Isl1* (Fig. S4a-c; 35) we obtained *Nkx2.1^loxP/loxP^*. *Isl1^+/cre^* mice, that we named Isl1*Nkx2.1*KO. Analysis of Isl1*Nkx2.1*KO embryos at E11.5 showed an almost complete absence of arcuate POMC positive-cells in contrast to control *Nkx2.1^loxP/loxP^* littermates (Fig. 7 a,b). Interestingly, the hypothalamic pattern of *Nkx2.1* expression is only altered in the mantle zone of the developing arcuate nucleus where most ISL1 positive-cells reside (Fig. 7 c-h). As observed in *Nkx2.1*KO@E9.5 embryos (Fig. 5i,l), Isl1*Nkx2.1*KO showed normal Isl1 levels (Fig. 7 e,f). Expression analysis of the proneuronal markers Ascl1 and neurogenin-3 in the developing hypothalamus showed no differences between control and Isl1*Nkx2.1*KO E11.5 embryos (Fig. 7i-l), suggesting that the lack of Nkx2.1 in ISL1 neurons does not affect neurogenesis in this brain region. Similarly, E12.5 Isl1*Nkx2.1*KO embryos showed a great reduction in the number of POMC neurons (Fig. S4 d-g). Thus, the lack of *Nkx2.1* expression specifically in ISL1 neurons impairs hypothalamic *Pomc* expression and confirms the critical role of Nkx2.1 in the establishment of melanocortin neurons.

**Figure 6.**
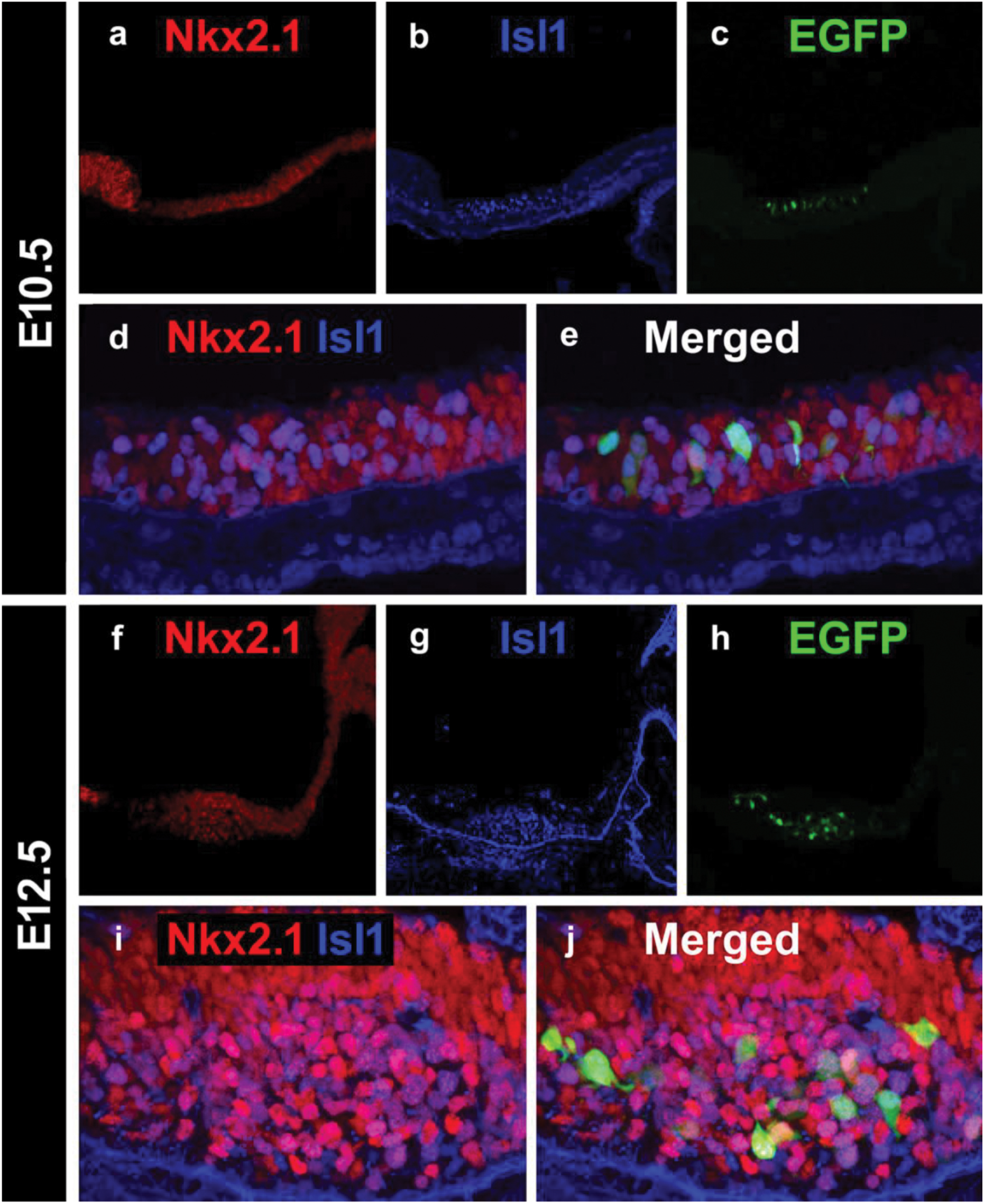
Overlapping expression patterns of *Nkx2.1*, *Isl1* and *Pomc* in the developing hypothalamus. Immunofluorescence analysis using anti-NKX2.1 (red) and anti-ISL1 (blue) antibodies in sagittal cryosections of E10.5 (a-e) and E12.5 (f-j) *Pomc*-EGFP mouse embryos. Higher magnification confocal images showing superimposed NKX2.1 and ISL1 immunopositive neurons (d and i) and triple merged with *Pomc*-EGFP immunoreactive neurons (e and j).

### Selective ablation of *Nkx2.1* from POMC neurons impairs *Pomc* expression and increases body weight

To investigate whether NKX2.1 participates in arcuate *Pomc* expression specifically in POMC neurons, we crossed *Nkx2.1^loxP/loxP^* mice with a BAC *Pomc*-Cre transgenic line (36). In the resulting Pomc*Nkx2.1*KO mice, the *Nkx2.1* alleles are inactivated in POMC neurons once *Pomc-Cre* expression begins. Immunofluorescence analyses of Pomc*Nkx2.1*KO embryos at E12.5 and E15.5 showed that the number and location of POMC neurons in the developing arcuate nucleus were identical to those observed in control littermates (Fig. 8 a-d), in contrast to what we found when *Nkx2.1* was ablated at earlier time points (Fig. 5 and 7). Given that *Nkx2.1* and *Pomc* also coexpress in the arcuate nucleus during postnatal life, we decided to evaluate whether *Nkx2*.1 plays a role in the transcriptional regulation of *Pomc* in adult mice. We found that the number of POMC neurons in the adult arcuate nucleus was normal, as assessed in 20-week-old *Pomc*-EGFP.Pomc*Nkx2.1*KO compound mice (Fig. 8 e,f). However, a quantitative RT-PCR study showed that hypothalamic *Pomc* mRNA levels from 20-week-old Pomc*Nkx2.1*KO mice are 38 % lower than those found in *Nkx2.1^loxP/loxP^* control littermates (Fig 9a). In agreement with the reduction in *Pomc* mRNA levels, Pomc*Nkx2.1*KO mice are heavier than their control siblings along the entire 5 to 19 week-old interval measured (Fig. 9b), although they displayed no differences in body length. Interestingly, Pomc*Nkx2.1*KO 19-week-old male mice displayed overweight, being 11% heavier than their control littermates (control: 32.5 +/− 0.8 g and Pomc*Nkx2.1*KO: 36.2 +/− 1.0 g; P<0.01). In addition, the livers and retroperitoneal fat pads of Pomc*Nkx2.1*KO mice were 16 % (Fig. 9c) and 120 % (Fig. 9d) heavier than those from control littermates, whereas other fat pads showed no statistically significant difference (Fig. 9d). Overall, our results demonstrate that NKX2.1 plays a fundamental cell-autonomous role in attaining normal *Pomc* expression levels in the adult arcuate nucleus and, consequently, in the regulation of normal body weight homeostasis.

**Figure 7.**
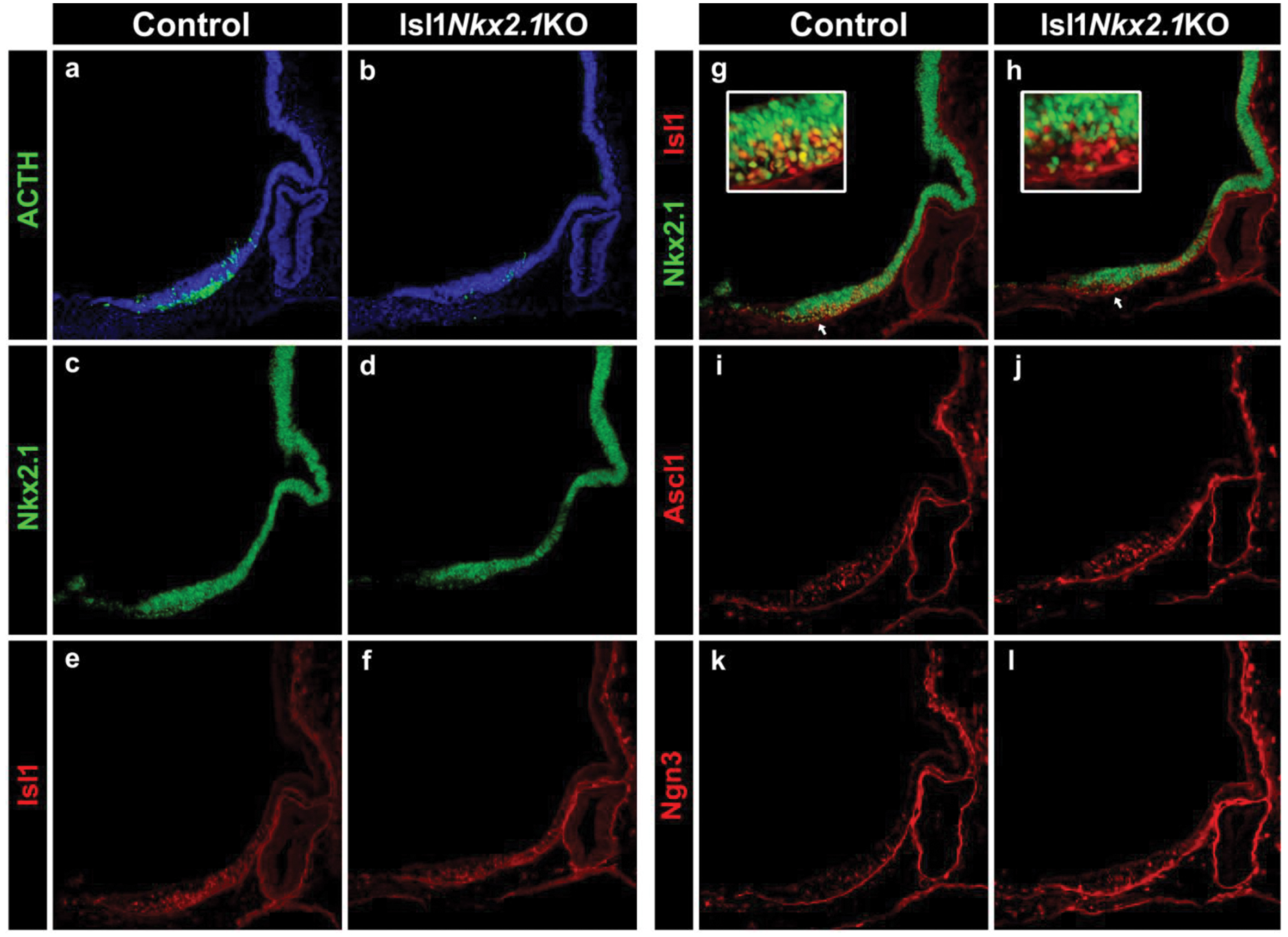
Specific deletion of *Nkx2.1* from ISL1+ neurons impairs the onset of *Pomc* expression. Immunofluorescence analysis in sagittal cryosections of E11.5 embryos using anti-ACTH (a,b), anti-NKX2.1 (c,d), anti-Isl1 (e,f), anti-Ascl1 (I,j) and anti-Ngn3 (k,l) antibodies. Loss of Nkx2.1 in Isl1+ neurons is evident by double labeling (g,h). Arrows point to the mantle zone of the mediobasal hypothalamus detailed in the squares above. DAPI is shown in blue.

**Figure 8.**
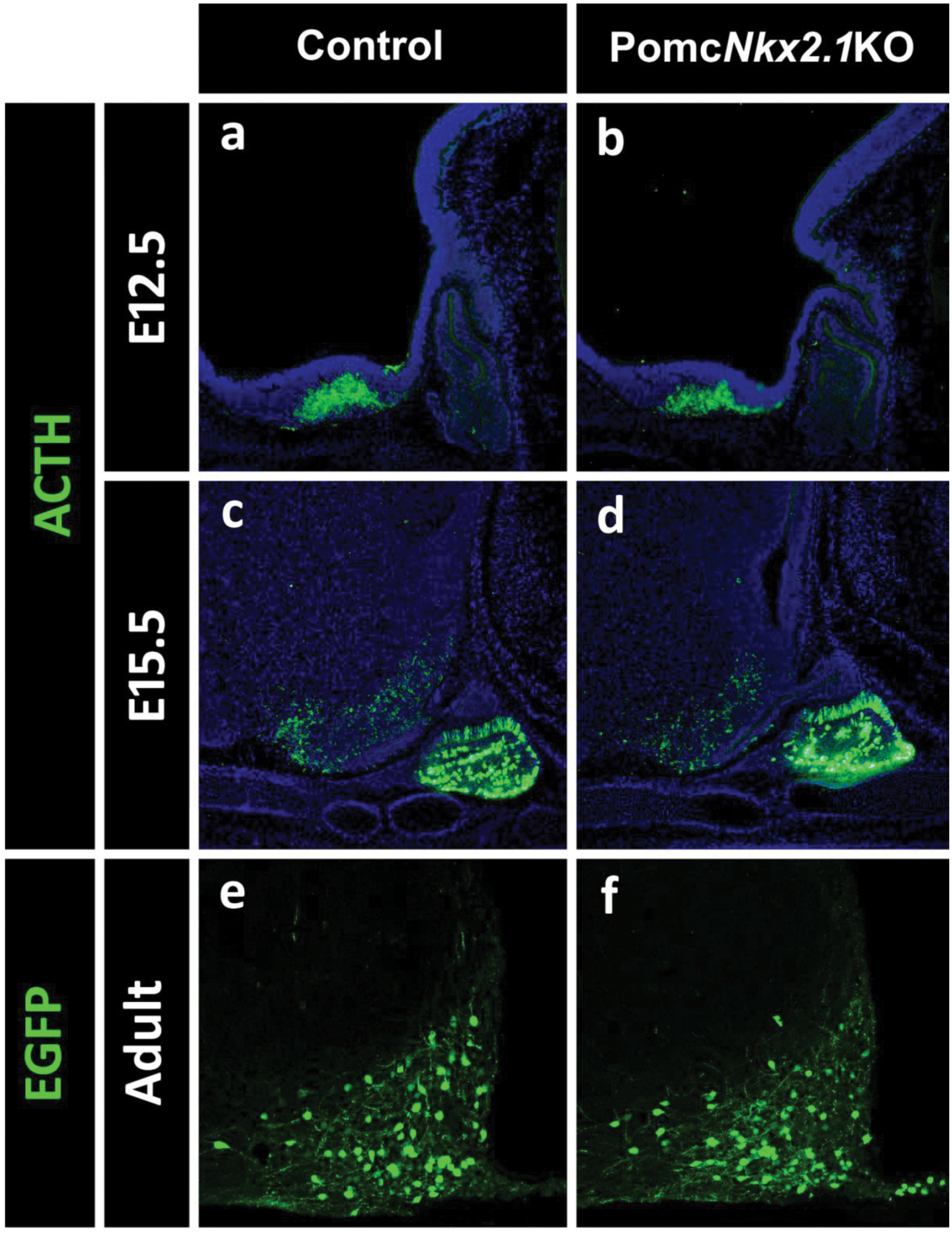
*Nkx2.1*-specific deletion in POMC neurons. (a-d) The number of ACTH+ cells in Pomc*Nkx2.1*KO mice is normal at E12.5 (a,b) and E15.5 (c,d) in the hypothalamus and pituitary. The number of POMC neurons in the adult arcuate nucleus of Pomc*Nkx2.1*KO mice is normal, as assessed in *Pomc*-EGFP.Pomc*Nkx2.1*KO mice (e,f).

**Figure 9.**
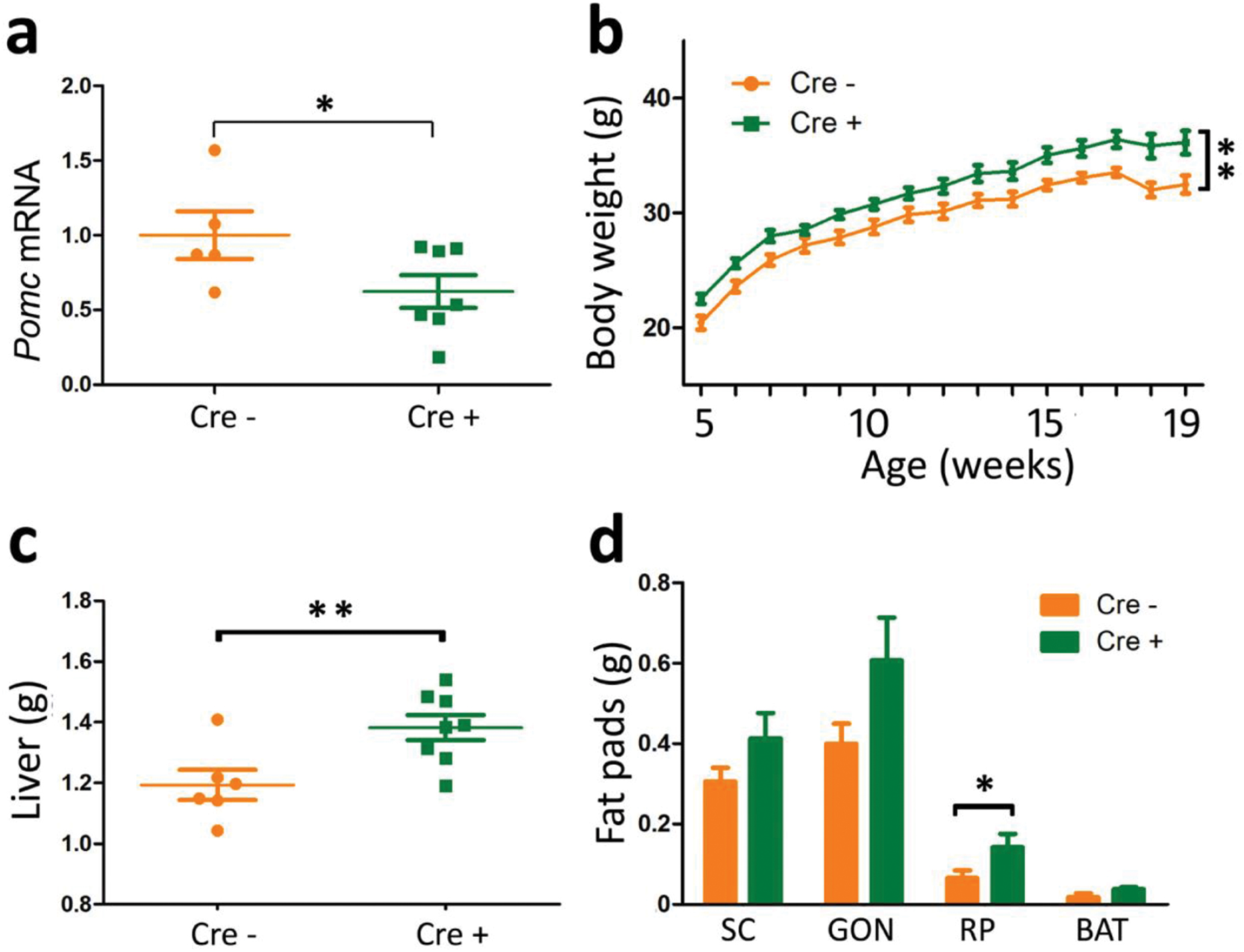
*Nkx2.1*-specific deletion in POMC neurons impairs *Pomc* expression and causes early onset obesity. (a) Quantitative RT-PCR shows reduced *Pomc* mRNA levels in the hypothalamus of adult Pomc*Nkx2.1*KO mice. (b) Body weight curves reveal that Pomc*Nkxc2.1*KO male mice are overweight, display heavier livers (c) and increased adiposity in the retroperitoneal fat pads (d). (a,c,d) Values represent the mean ± SEM (n = 5–8). *P < 0.05, **P < 0.01 (Student’s t test). (b) Values represent the mean ± SEM (n = 6–8). *P < 0.01 (RMA).

## Discussion

In this study we combined molecular, genetic, cellular, and functional approaches to demonstrate that the homeodomain transcription factor NKX2.1 is necessary to activate and maintain arcuate-specific *Pomc* expression from early development to adulthood. Therefore, Nkx2.1 determines the identity of hypothalamic melanocortin neurons. Specifically, we show that: (i) the neuronal *Pomc* enhancers nPE1 and nPE2 contain canonical NKX binding sites which are highly conserved in all mammalian orders; (ii) *Pomc* coexpresses with *Nkx2.1* from its onset at E10.5 and throughout the entire lifespan in the arcuate nucleus; (iii) NKX2.1 binds *in vitro* to DNA fragments from nPE1 and nPE2 carrying NKX binding motifs and *in vivo* to nPE1 and nPE2 in hypothalamic chromatin extracts; (iv) the NKX binding motifs present in nPE1 and nPE2 are essential for their enhancer activity in arcuate neurons; (v) early expression of *Nkx2.1* in the developing ventral hypothalamus is necessary for the onset of hypothalamic *Pomc* expression, a first hallmark of postmitotic melanocortin neurons; (vi) *Nkx2.1* ablation from hypothalamic ISL1+ neurons prevents *Pomc* expression demonstrating that the sole presence of Isl1 is unable to activate *Pomc* and; (vii) *Nkx2.1* ablation from POMC neurons after E10.5 reduces *Pomc* mRNA levels and increases body weight and adiposity in adult mice.

*Nkx2.1* is expressed as early as E7.5 in the anterior portion of the mouse neural tube and hours later is found in neuronal progenitors of the hypothalamic anlage, where it specifies ventral lineages while repressing dorsal and alar fates (25, 26). Ablation of *Nkx2.1* at this early stage impairs the formation of the ventral hypothalamic primordium which undergoes dorsalization (25). In addition to this essential early role defining the ventral identity of the developing hypothalamus, *Nkx2.1* is later expressed in postmitotic cells, suggesting that it may also be involved in the differentiation and maintenance of mature cellular phenotypes in this brain region (27,37). In fact, in this study we have found that NKX2.1 plays a crucial role in establishing the early identity of melanocortin neurons by promoting the onset of *Pomc* expression in the ventral hypothalamus and also participates in the transcriptional regulation of *Pomc* until adulthood. Thus, NKX2.1 and its cognate NKX binding motifs present in the *Pomc* neuronal enhancers nPE1 and nPE2 contribute to sustain the supra threshold levels necessary to regulate a normal body weight and adiposity. Moreover, since *Pomc* mRNA and peptides are the archetypal and unique markers that define the neuronal melanocortinergic phenotype, the multiple developmental roles that NKX2.1 plays in the future ventral hypothalamus assure first, the formation of the tuberal portion of the mediobasal hypothalamus from where the arcuate nucleus originates; second, the determination of the identity of POMC neurons in this region and; third, the maintenance of high expression levels of *Pomc* to regulate body weight.

Our results shed light on the multistep genetic program that NKX2.1 initiates in the anterior tip of the neural tube during the early developmental stages of the anterior brain, and that ends up with the integration of fully functional neuronal circuits including those controlling food intake and energy balance. This complex program, strongly dependent on NKX2.1, includes the morphogenesis of the arcuate nucleus followed by the birth and maturation of arcuate neurons expressing *Pomc*. The execution of this elaborate developmental program has been recently mimicked *in vitro* using, at starting points, human embryonic stem cells or induced pluripotent stem cells (38, 39). Upon treatment with the morphogen Sonic Hedgehog Homolog and inhibitors of the SMAD and Notch pathways these cells acquire features of ventral hypothalamic neural progenitors including the expression of *Nkx2.1* (39). Further inhibition of Notch signaling differentiates NKX2.1 positive cells to a population of neuronal phenotypes typical of the arcuate nucleus including POMC, NPY/Agrp, somatostatin and TH/dopamine (39).

In addition to its critical role in the developmental plan of the ventral hypothalamus and telencephalic medial ganglionic eminence (26), NKX2.1 drives the differentiation of particular cell types in peripheral organs such as the thyroid gland (32) and the lungs (40). As we found in hypothalamic POMC neurons, NKX2.1 participates not only in the morphogenesis of the thyroid gland but also in the differentiation of thyrocytes and in the regulation of the expression of thyroid-specific genes such as thyroglobulin (41). The concomitant function of NKX2.1 in these three organs is evident in a familial clinical condition known as brain-lung-thyroid syndrome, characterized by congenital hypothyroidism, infant respiratory distress and benign hereditary chorea, which is found in patients carrying heterozygous missense or nonsense mutations in the coding region of human *NKX2.1* (41). Some *NKX2.1* mutations involve two or just one of these conditions and, in addition, familial cases of ataxia and pituitary abnormalities have been reported (42). Although none of these cases reported comorbidity with overweight or obesity, our observation that *Nkx2.1* deficiency in POMC neurons induces increased adiposity and overweight in the mouse may be of biomedical relevance since polymorphisms leading to very low levels of NKX2.1 in the ventral hypothalamus or in NKX binding motifs in neuronal *POMC* enhancers may reduce *POMC* expression and consequently impair the control of food intake and energy balance. Innumerable genome-wide association studies have been performed during the last decades identifying more than 100 different loci potentially associated with high body mass index, type II diabetes, increased adiposity or high leptin levels (43-45). Although the individual contribution of the vast majority of these variants is quantitatively irrelevant, a polymorphic sequence present in a conserved intronic region of the FTO gene gained special significance because its statistical power has been replicated in other unrelated studies (46, 47). Further genetic studies showed that this conserved intronic sequence is a tissue-specific enhancer that controls not FTO but a distant gene coding for the TF Irx3 (48). Another exceptional single locus is *POMC*, since a number of genome-wide studies have found highly significant linkage scores between obesity-related traits and a genomic segment in chromosome 2 near *POMC* (49, 50)(51). Although polymorphisms in *POMC* coding sequences do not appear to account for these associations (52), it is likely that mutations in noncoding regulatory elements may alter *POMC* transcript levels and modify the relative amount of central melanocortins.

Our finding that NKX2.1 binds to canonical NKX binding motifs present in nPE1 and nPE2 adds to the adaptive partial redundancy of arcuate *Pomc* expression that relies on the presence of these two enhancers for full transcriptional activity (20). During mammalian evolution, nPE1 and nPE2 were independently exapted (co-opted) from different types of retroposons (19). This long lasting evolutionary process in which two different retroposon-derived sequences became functional neuronal *Pomc* enhancers involved the independent acquisition of NKX binding sites in each of them, as we previously found for ISL1 binding sites (21). Because the individual ablation of *Nkx2.1* or *Isl1* from early developmental stages prevents the onset of hypothalamic Pomc expression we conclude that the combinatorial presence of NKX2.1 and ISL1 is necessary to determine the identity of arcuate melanocortin neurons. In fact, in this study we show that *Pomc* is expressed in NKX2.1+, ISL1+ neurons. However, since the expression territories of these two TFs in the arcuate nucleus is much broader than that of *Pomc*, it is clear that these two TFs are not sufficient for arcuate-specific *Pomc* expression and that at least another TF, yet to be discovered, is necessary to dictate the final identity of POMC neurons.

## Acknowledgments

The authors thank Marta Treimun for excellent technical assistance in mice production, Dr. Shioko Kimura for kindly providing the *Nkx2.1* conditional mutant mice and Lucía Franchini for valuable protocols and ideas. This work was supported by National Institutes of Health Grant R01 DK068400 (to M.J.L. and M.R.), Agencia Nacional de Promoción Científica y Tecnológica (to M.R.) and UBACYT (M.R.).

## Author contributions

D.P.O, M.B.T., F.S.J.d.S., M.J.L., and M.R. designed research; D.P.O, M.B.T., F.S.J.d.S., S.N., and M.R. performed research; M.J.L. and M.R. contributed new reagents and analytic tools; D.P.O, M.B.T., F.S.J.d.S and M.R. analyzed data and prepared figures; M.R. wrote the paper. D.P.O, M.B.T., F.S.J.d.S., S.N., M.J.L., and M.R revised the paper.

---

**Fig. S1. nPE1 multiple sequence alignment.** ClustalW of nPE1 sequences from a representative variety of eutherian (placental) mammalian species. The unique canonical NKX-binding site present in this enhancer is labeled in a blue box.

**Fig. S2. nPE2 multiple sequence alignment.** ClustalW of nPE2 from a representative variety of mammalian species. The species include placental mammals as well as one marsupial (opossum, *Monodelphis domestica*) and a monotreme (platypus, *Ornithorhynchus anatinus*). The two NKX-binding sites present in nPE2 are shown within blue squares.

**Fig. S3. Pomc-EGFP transgenic mice express the reporter fluorescent marker in all POMC neurons.** ACTH and EGFP coexpress in the developing mediobasal hypothalamus at E10.5 and E12.5 as shown by immunofluorescence.

**Fig. S4. The *Isl1^+/cre^* mouse model.** (a-c) Immunofluorescence using an anti-ACTH antibody shows that the number of POMC neurons in Isl1*Nkx2.1*KO E12.5 embryos is greatly reduced in comparison to *Isl1^+/+^* and *Isl1^+/cre^* controls. Although *Isl1^+/cre^* mice are haploinsufficient for *Isl1*, they displayed a normal number of POMC neurons in agreement with our previous analysis of *Isl1^+/−^* mice (21). (d-g) Isl1*Nkx2.1*KO and control E12.5 mouse embryos. Immunofluorescence using an anti-ACTH (d,e) or anti-Nkx2.1 (f,g) antibodies. The white arrow points to the mantle zone of the future arcuate nucleus, where NKX2.1 disappears in Isl1*Nkx2.1*KO embryos.

## Materials and Methods

### Breeding of Mice

Mice were housed in ventilated cages under controlled temperature and photoperiod (12-h light/12-h dark cycle, lights on from 7:00 AM to 7:00 PM), and fed *ad libitum.* All procedures followed the Guide for the Care and Use of Laboratory Animals and in agreement with the INGEBI-CONICET institutional animal care and use committee. *Nkx2.1^loxP/loxP^* mice were provided by Dr. Shioko Kimura and maintained as homozygotes on a C57BL6/J background. CAAG-CreERT mice (1) were obtained from the Jackson Laboratory (B6.Cg-Tg[CAG-Cre/Esr1]5Amc/J) and were intercrossed with *Nkx2.1^loxP/loxP^* mice to generate Ind*Nkx2.1*KO strain. *Isl1*-Cre (2) mice were obtained from Jackson Laboratory (*Isl1^tm1(cre)Sev^/*J) and maintained as heterozygotes on a C57BL6/J background. Genotyping was performed by PCR using genomic DNA extracted from ear biopsies or embryo fragments. All primer sequences used are available in Table S1.

### Transgene and Transgenic mice production

Transgene nPE2*Pomc*-EGFP was previously reported (3,4). Transgene nPE2(NKX*)*Pomc*-EGFP is identical except for carrying two transversion mutations in each of the two NKX binding sites present in nPE2 (TC<u>AA</u>G→TC<u>CC</u>G and TC<u>AA</u>T→TC<u>CC</u>T). This transgene was generated using a standard megaprimer PCR protocol as previously described (5) but with the primers described in Table S1. Introduction of mutations was confirmed by sequencing. Transgenes were digested with *NotI* and *SalI* restriction enzymes and the fragment of interest was separated and extracted from a 0.8% agarose-TBE gel with QIAquick Gel Extraction Kit (Qiagen, Cat No.: 28704). The transgene was further concentrated and purified with DNA Clean & ConcentratorTM-5 (Zymo Research, Cat No.: D4014) and eluted in 5 ul EmbryoMax injection buffer (Millipore, Cat No.: MR-095-10F). Transgenic mice were produced by pronuclear microinjection of the transgene in Fvb zygotes. Transgenic pups were identified by PCR using “M329” and “M330” primers. nPE2 was amplified using “Δ2.3” and “Δ2.5” primers cloned using pGEM-T Easy Vector System I (Promega, Cat. No.: A1360) and sequenced to confirm mutations. Founder mice were intercrossed with Fvb *wild type* mice and F1 newborns were analyzed.

### Knockout mice generation

CRISPR/Cas9 system was used to generate deletions encompassing NKX binding site in nPE1. A CRISPR guide was selected using crispr.mit.edu website of Zhang Lab. Guide CCCCACAATGGGGCTTGAAG was selected because it falls next to *Nxk2.1* binding site and had a high quality score and thus low chances of producing off-target mutations. The guide was sub-cloned in plasmid DR274 (Addgene Plasmid #42250) and sgRNA was synthesized using MEGAshortscript T7 Transcription Kit (Ambion, Cat# AM1354). Cas9 mRNA was synthesized from plasmid MLM3613 (Addgene Plasmid #42251) using mMESSAGE mMACHINE™ T7 Transcription Kit (Ambion, AM1344) and Poly(A) Tailing Kit (Ambion, AM1350). Cas9 mRNA (50 ng/ul) and sgRNA (50 ng/ul) were injected in the cytoplasm of *Pomc^+/^*^ΔnPE2^ C57BL6/J zygotes. Founder mice were analyzed by PCR using primers available in Table S1 and PCR products subcloned in pGEM.T Easy Vector for sequencing. A strain carrying a 34 bp deletion (chr12:3,942,178-3,942,211) in nPE1 in a ΔnPE2 allele was selected and maintained in a C57BL6/J background and F2 mice were analyzed.

### Tamoxifen injections

Tamoxifen (T5648, Sigma) was dissolved in sesame oil (S3547, Sigma) at 15 mg/mL by sonication. 1 ml aliquots were stored at −20°C and thawed for 10 minutes at 37°C before injection. Pregnant females were injected intraperitoneally with a dose of 133 mg/kg.

### Tissue collection and embedding

Pregnant mice at 10.5-15.5 days post coitum (dpc, the day of detection of vaginal plug was considered dpc 0.5) were dislocated and embryos were collected and washed in cold RNAse-free PBS. Fixation was performed with 4% PFA-PBS at 4°C, for a period of time dependent on developmental stage. 10.5, 11.5 and 12.5 dpc embryos were fixed for 2hs, 15.5 dpc embryos’ heads were dissected and fixed for 4hs and newborns’ heads were fixed overnight. Samples were then washed in cold PBS and cryoprotected with 10% sucrose-PBS overnight. Tails or posterior limbs were cut in order to obtain tissue for genotyping. Tissues were then stabilised in 10% sucrose-10% gelatin-PBS at 37°C for 30 min prior to freezing. Adult animals were perfused using 4% PFA and brains were collected and fixed overnight in 4% PFA at 4°C. Whole brains were then washed in cold PBS and cryoprotected with 10% sucrose-PBS overnight. Hypothalamic sections were then cut grossly using a metallic adult brain mold prior to freezing. Gelatin blocks containing the tissues were stuck on cork sheets with Tissue-Tek OCT compound (Sakura Finetek, CAt. No.: 4583). Gelatin blocks were snap-frozen in 2-Methylbutane (Sigma, Cat No.: M32631) at −55°C and stored at −80°C for up to 1 year. 20 μm slices were cut in a cryostat (Leica, CM 1850) and mounted on microscope slides (Fisherbrand Superfrost Plus, Cat No.: 12-550-15). Sections were then air-dried overnight and stored at −20°C until use.

### Immunofluorescence

Immunofluorescence was performed as described (5). Primary antibodies used were: rabbit anti-rACTH (1:1000, National Hormone and Peptide Program); mouse anti-ISL1 (1:20, 40.2D6, Developmental Studies Hybridoma Bank, University of Iowa); rabbit anti-NKX2.1 (1:1000, ab40880, Abcam); rabbit anti-NKX2.1 (1:1000, sc-13040, Santa Cruz Biotechnology); chicken anti EGFP (1:2000, GFP-1020, Aves Lab); mouse anti Ascl1 (1:200, 556604, BD Pharmingen); mouse anti Ngn3 (1:2000, F25A1B3, Developmental Studies Hybridoma Bank, University of Iowa). Secondary antibodies were either anti-mouse or anti-rabbit AlexaFluor 555 (1:1000, Invitrogen); anti-mouse, anti-rabbit AlexaFluor 488 (1:1000, Invitrogen) or anti-chicken Alexa Fluor 488 (1:2000, Jackson Immuno Research, Cat. No.: 703-545-155). Depending on the marker antigen, immunohistochemistry experiments were repeated in 2 to 5 different embryos, with consistent results. Nuclei were stained with a 1 mg/l dilution of DAPI for 10 minutes and then washed twice before mounting with V Confocal images for co-expression assays and cell counting were obtained using a FV300 system combined with a BX51 microscope (Olympus) or Leica Confocal TCS-SPE. The number of EGFP+ or fluorescently-immunolabeled ACTH^+^ or ISL1^+^ cells were counted by confocal microscopy. Total number of labelled cells per animal are derived from 8 slices spanning the whole hypothalamus of coronally cut adult brains (*Pomc*-EGFP) or 4-5 slices spanning the whole hypothalamus of sagittal cut embryos (ACTH^+^ and ISL1^+^).

### *Pomc* mRNA Quantification by RT-qPCR

Whole mouse adult hypothalami or embryo heads were dissected, collected in ice-cold TriPure Isolation Reagent (Sigma, Cat. Nr. 11667165001) and stored at −80°C until RNA extraction, which was performed following manufacturer’s instructions. RNA integrity was assessed by gel electrophoresis; clear 28S and 18s rRNA were observed in an approximate 2:1 ratio. Quantification was performed using a Nanodrop and 260/280 and 260/230 ratios were checked to assess purity. 1 μg of RNA was DNAse I (Ambion Cat. No.: AM2222) treated and used for first-strand cDNA synthesis, using High Capacity Reverse Transcription Kit with random primers (Applied Biosystems, Cat Nr. 4368814). Primers were designed with Primer 3 program. *Pomc* mRNA was quantified using primers spanning exons 2 and 3 relative to internal control genes β2-microglobulin (B2m) and β-actin (Actb). Primer sequences are listed in Table S1. Samples were run in triplicate in an Applied Biosystems 7500 Real-Time PCR System machine using Power Up SYBR Green Master Mix (Applied Biosystems, Cat. No.: A25472). Melt curves were analyzed to confirm specificity of PCR product. Relative quantification was done by interpolating Ct values in standard curves or by 2-ΔΔCT.

### Electrophoretic mobility shift assay (EMSA)

A mouse *Nkx2.1* cDNA clone (ATCC, No.: 10698306) was subcloned into the expression vector pGEX 4T3 using NcoI and EcoRI restriction enzymes and transformed into Escherichia coli BL21 strain. Protocols for obtaining bacterial extracts, radioactive labeling of DNA probes and preparation of EMSA reactions were previously described (6). After electrophoresis, gels were dried and exposed to X-ray film (Kodak BioMax MS) or Storage Phosphor Screen that was subsequently scanned in a STORM 860 Phospho Imager using ImageQuant5.2 software (Amersham Biosciences). Probe sequences are shown in Table S1.

### Chromatin Immunoprecipitation (ChIP)

Hypothalami from 5 adult mice were collected and fixed in 1% PFA-PBS for 20 minutes at 4°C. Samples were sonicated to obtain 200-400bp chromatin fragments. 40ug of chromatin were immunoprecipitated with 5 ug of NKX2.1 antibody (sc-13040x, Santa Cruz Biotechnologies) or Normal Rabbit IgG (12-370, Merck Millipore). After overnight incubation, antibody-chromatin complexes were pulled down using a mixture of salmon sperm DNA blocked Protein G PLUS-agarose and Protein A-agarose (sc-2002, sc-2001, Santa Cruz Biotechnologies). Beads were washed with Low salt (Tris-HCl pH:8 10mM, Triton X-100 1%, EDTA 2mM, SDS 0.1%, NaCl 150mM), High Salt (As Low salt but with NaCl 450mM), LiCl (Tris-HCl pH:8 10 mM, LiCl 0.25M, IGEPAL-CA630 1%, EDTA 1mM, Na deoxycholate 1% (w/v)) and TE (twice) buffers. Elution was performed overnight at 65°C with 10ug of Proteinase K in Tris-HCl pH:8 10mM, SDS 1%, NaCl 340mM. DNA was PIC-IC extracted and precipitated with sodium acetate and ethanol. Immunoprecipitated fragments were analyzed by end-point PCR (Primer information is available in Table 1).

### Statistics

All data presented are the mean ± SEM and were analyzed using GraphPad Prism by repeated measures ANOVA or Student’s t test unless otherwise stated. In the case of RT-qPCR performed in embryos, variances were unequal as determined by Bartlett’s test and dependent on the mean. In order to perform ANOVA, variance was modelled using VarPower function of “nmle” package in R Studio (7). Post hoc pairwise comparisons between groups were performed by Tukey test. P values less than 0.05 were considered significant.

---

**Table S1.**
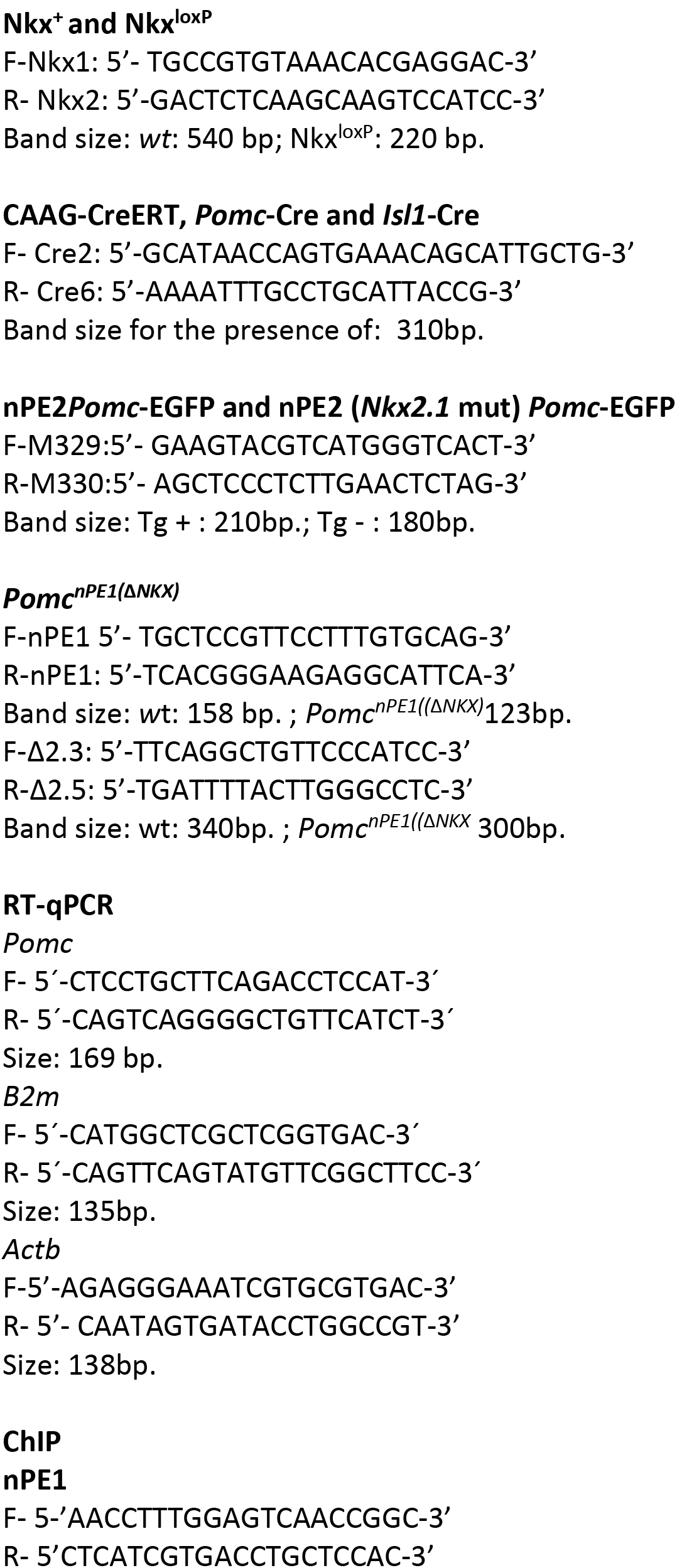
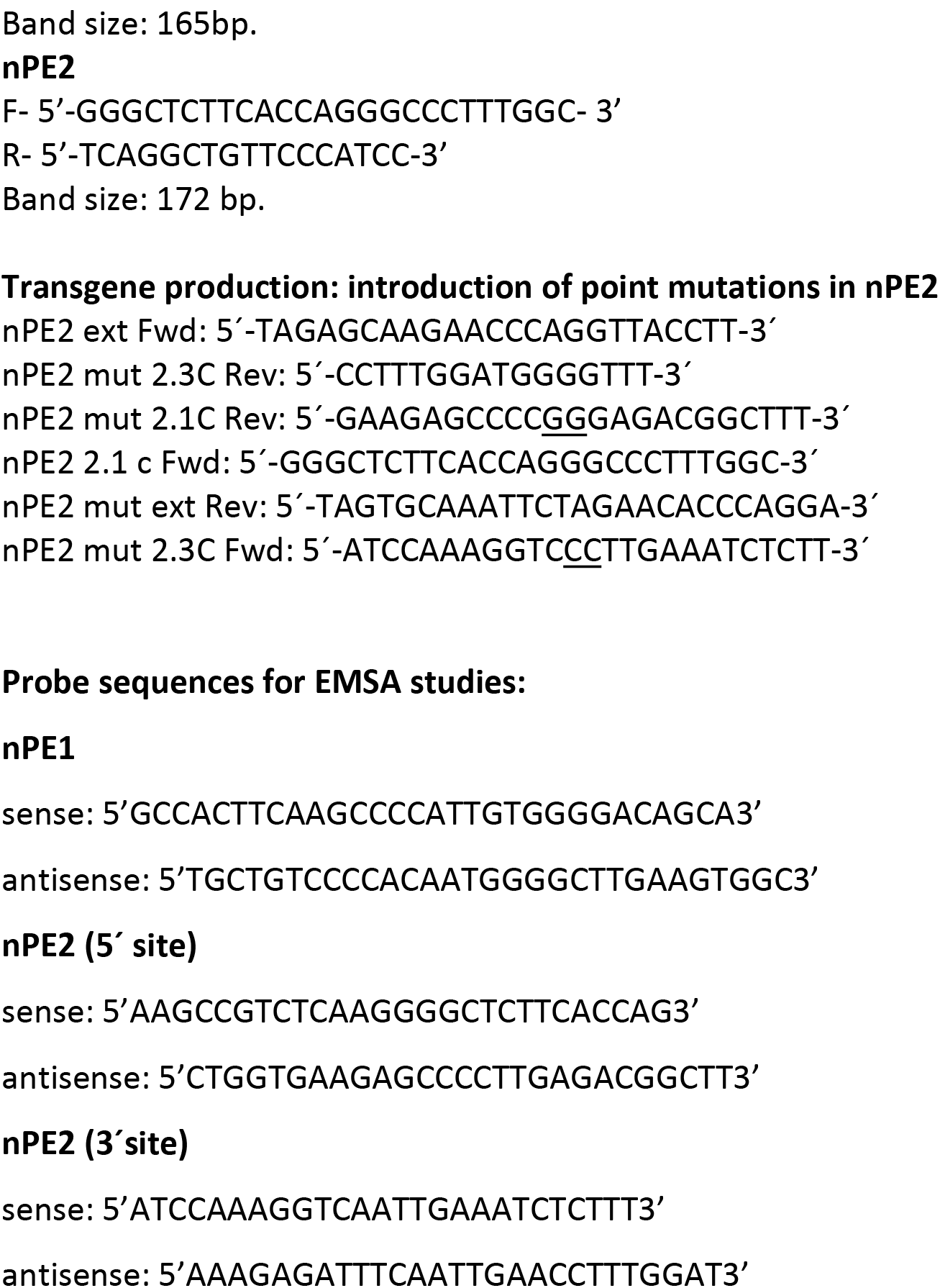
Primer sequences used to detect different alleles and mutations and probes for EMSA studies.

